# Neurotransmitter release is triggered by a calcium-induced rearrangement in the Synaptotagmin-1/SNARE complex primary interface

**DOI:** 10.1101/2024.06.17.599435

**Authors:** Estelle Toulmé, Andrea Salazar Lázaro, Thorsten Trimbuch, Josep Rizo, Christian Rosenmund

## Abstract

The Ca^2+^ sensor synaptotagmin-1 triggers neurotransmitter release together with the neuronal SNARE complex formed by syntaxin-1, SNAP25 and synaptobrevin. Moreover, synaptotagmin-1 increases synaptic vesicle priming and impairs spontaneous vesicle release. The synaptotagmin-1 C_2_B domain binds to the SNARE complex through a primary interface via two regions (I and II), but how exactly this interface mediates distinct functions of synaptotagmin-1, and the mechanism underlying Ca^2+^-triggering of release is unknown. Using mutagenesis and electrophysiological experiments, we show that region II is functionally and spatially subdivided: binding of C2B domain arginines to SNAP-25 acidic residues at one face of region II is crucial for Ca^2+^-evoked release but not for vesicle priming or clamping of spontaneous release, whereas other SNAP-25 and syntaxin-1 acidic residues at the other face mediate priming and clamping of spontaneous release but not evoked release. Mutations that disrupt region I impair the priming and clamping functions of synaptotagmin-1 while, strikingly, mutations that enhance binding through this region increase vesicle priming and clamping of spontaneous release, but strongly inhibit evoked release and vesicle fusogenicity. These results support previous findings that the primary interface mediates the functions of synaptotagmin-1 in vesicle priming and clamping of spontaneous release, and, importantly, show that Ca^2+^-triggering of release requires a rearrangement of the primary interface involving dissociation of region I, while region II remains bound. Together with modeling and biophysical studies presented in the accompanying paper, our data suggest a model whereby this rearrangement pulls the SNARE complex to facilitate fast synaptic vesicle fusion.

**Significance statement:** The synaptic SNARE complex and synaptotagmin-1 are required for fast neurotransmitter release. The functions of synaptotagmin-1 in preparing synaptic vesicles for fusion and executing the triggering step have been proposed to be regulated through interactions with the SNARE complex via the so-called primary interface. Using site-directed mutagenesis and functional analysis in neurons, we now show that synaptotagmin-1 mediates its release preparatory functions via two contact sites with the SNARE complex at this interface. During Ca^2+^ triggering, synaptotagmin-1 continues to contact the SNAREs at one site but disconnects the other site. We propose that this switch generates a pulling force on the SNARE complex that in turn triggers release. Biochemical and modeling studies described in the accompanying paper support this hypothesis.

## Introduction

Neuronal communication relies on the release of neurotransmitters by Ca^2+^-triggered synaptic vesicle exocytosis, which occurs rapidly (< 1 ms) after Ca^2+^ influx into the presynaptic terminal. This fast speed is possible because release is controlled by a sophisticated protein apparatus that first tethers synaptic vesicles (SVs) to the plasma membrane, then docks-primes SVs in a release-ready state and mediates very fast membrane fusion upon Ca^2+^ influx (1). At the core of this apparatus is the complex formed by the vesicular soluble N-ethylmaleimide sensitive factor attachment protein receptor (SNARE) protein synaptobrevin, and the plasma membrane SNAREs, syntaxin-1 and SNAP25, which form a trans-SNARE complex that consists of a four-helix bundle and brings the SV and the plasma membrane together (2). SV fusion is thought to be triggered by extension (zippering) of the synaptobrevin and syntaxin-1 helices to the juxtamembrane (jxt) linkers that precede their transmembrane (TM) regions (3–5), which facilitates encounters of the hydrophobic lipid acyl chains of both membranes at the polar membrane interface (6). Fast, synchronous neurotransmitter release also requires the SV protein synaptotagmin-1 (Syt1), which is believed to have multiple roles: First, it promotes the supply of fusion competent vesicles by boosting SV docking and priming (7). Second, it increases signal-to-noise properties at synapses by clamping spontaneous release (8). Third, it is the Ca^2+^ sensor that triggers action potential-evoked release (9–11).

Only the third function is Ca^2+^ dependent. The Ca^2+^ triggering step involves the binding of the Ca^2+^-binding loops of the two C_2_ domains of Syt1 (C_2_A and C_2_B) to phospholipid membranes (9, 11). The Syt1 C_2_B domain also contains two patches of basic residues that are distantly located from the Ca^2+^ binding region and that are critical both for its vesicle docking/priming function and for its role in Ca^2+^-triggered release (7, 12–15). One of these patches, known as the polybasic region, binds to acidic phospholipids, including phosphatidylinositol 4,5-bisphosphate (PIP_2_), and is generally believed to bind to the plasma membrane (16, 17). The other basic patch includes three arginine residues (R281, R398 and R399) that bind to the SNARE complex and form part of a so-called primary interface between the C_2_B domain of Syt1 and the SNARE complex revealed by X-ray crystallography (15). The primary interface is formed mostly by two regions, one involving these three arginines of the C_2_B domain as well as acidic residues from syntaxin-1 and SNAP25 (referred to as region II), and another region involving E295 and Y338 of the C_2_B domain (region I) (Fig. S1A,B). Functional and biochemical experiments have suggested that all three of the proposed functions of Syt1 occur through or are modulated by primary interface interactions. Mutations in SNAP25 residues that participate in either region I (D166/E170) or region II (D51/E52/E55) impair evoked responses (18). The region I mutants also display reductions in vesicle priming and enhancements in spontaneous release, while mutations in region II enhance spontaneous release without affecting RRP size (18). Mutation of R398 and R399 of the Syt1 C_2_B domain at region II of the primary interface, which abolishes binding through this interface (17), severely disrupts evoked release and vesicle priming (7, 13, 15, 19). Intriguingly, a E295A/Y338W mutation in region I impaired evoked release but rescued spontaneous release clamping (15) and enhanced binding of the C_2_B domain to the SNARE complex (17). This finding led to the proposal that the primary interface is important for priming but is dissociated upon Ca^2+^ binding to Syt1 to enable evoked release (17).

While these data support an overall role of the primary interface in the regulation of Syt1 function, the described phenotypic patterns are complex, and a systematic assessment of how the primary interface contributes to the three functions of Syt1 is lacking. Moreover, the exact mechanism for how Syt1 is coupled to SNARE-mediated membrane fusion is unknown.

In this study, we aimed to tease apart the roles of the Syt1-SNARE primary interface in the pre-Ca^2+^-triggering functions of SV priming and spontaneous fusion clamping, as well as in Ca^2+^-triggering of release. With this goal, we performed a systematic structure-function analysis of the primary interface between Syt1 and the SNARE complex, including mutations in the Syt1 C_2_B domain as well as syntaxin-1 and SNAP25. We find that primary interface interactions mediate the functions of Syt1 in SV priming, clamping of spontaneous release, and Ca^2+^-triggered release, all in full agreement with previous studies (7, 15, 18). Moreover, we demonstrate that, while Syt1 loss of function mutations in region I disrupt all functions of Syt1, a gain of function mutation shows a unique phenotype: the Syt1 E295A mutation increases vesicle priming and enhances clamping of spontaneous release but impairs Ca^2+^-triggered release. In contrast, mutations in region II show a dual behavior, affecting either pre-Ca^2+^ or post-Ca^2+^ functions. These results and the results from the accompanying paper (20) strongly suggest that the interface between Syt1 and the SNARE complex undergoes a rearrangement that mediates the transition from a primed/clamped state of SVs to a state that selectively promotes Ca^2+^-triggered fusion. During this rearrangement, region I must dissociate, and region II is remodeled such that interactions supporting pre-Ca^2+^ triggering functions are relieved and interactions promoting Ca^2+^-triggered release are favored.

## Results

### Region II separates into two functionally distinct interfaces

To examine the functional roles of individual residues of the primary interface, we created a series of point mutants, guided in part by the results of previous mutational studies (7, 13, 15, 18) and by those of a screen for mutations that suppress the lethality caused by expression of Syt1 bearing mutations that abolish Ca^2+^-binding to the C_2_B domain in *Drosophila* (21). We generated single or double point mutants of Syt1 in region I (E295A, Y338D, Y338W, E295A/Y338W), in a small hydrophobic region between regions I and II (A402T), a combination of both (Y338D/A402T), and in region II (R281A, K288A, R281A/K288A, R398Q, R399Q, R398Q/R399Q). We also created region II mutations in syntaxin-1 (D231N/E234Q/E238Q) and SNAP25 (D51N, E52Q, E55Q, D51N/E52Q). The mutants were expressed as lentiviral rescues in the corresponding genetic knockout (KO) neurons (Syt1/7 DKO, SNAP25 KO, Syntaxin-1A/1B DKO). As a control, we used the corresponding rescue expression of the WT Syntaxin-1A, SNAP25 or Syt1. The efficacy of expression of all rescue constructs was monitored by Western blot analysis and compared to WT rescue expression (Fig. S2-S5). Individual hippocampal neurons were cultured on astrocytic micro-islands to avoid multi-neuron connectivity and to enable simultaneous and quantitative analysis of the three release functions, priming, spontaneous release and Ca^2+^-triggered release using electrophysiological experiments. Only glutamatergic neurons were incorporated in the analysis.

We first focused on acidic residues from the SNAREs in region II of the primary interface (Fig. 1A). Previous mutagenesis experiments on region II of SNAP25 showed that a triple D51A/E52A/E55A mutation disrupted Ca^2+^-triggered release but not vesicle priming(18). To investigate the individual roles of these acidic residues, we generated D51N, E52Q, and E55Q single mutants and a D51N/E52Q double mutant (Fig. 1A,B). Conversely, a study of region II of syntaxin-1 only mutated D231 among acidic residues in this region and found that a D231N mutation did not impair priming or evoked release but affected clamping (22). To examine the effects of mutating more than one residue in this region of syntaxin-1, we created E228Q/D231N and D231N/E234Q/E238Q mutants (Fig. 1J). Evoked excitatory postsynaptic currents (EPSCs) for the SNAP25 E52Q mutant as well as for the syntaxin-1 E228Q/D231N and D231N/E234Q/E238Q mutant expressing neurons were comparative in size with WT-rescue neurons (Fig. 1C,D,K,L). In contrast, the SNAP25 D51N and E55Q single mutants as well as the D51N/E52Q double mutant displayed a severe reduction in evoked release (Fig. 1C,D,K,L). Next, synaptic vesicle priming function was assessed by quantifying the readily releasable vesicle pool (RRP) size using pulsed application of hypertonic solutions (23). We found that the RRP size was normal for all SNAP25 mutants tested (Fig. 1E,F) but was reduced in the syntaxin-1 E228Q/D231N double and D231N/E234Q/E238Q triple mutants, (Fig. 1M,N). By taking RRP size and evoked responses into account together, we calculated the vesicular release probability (Pvr) and observed that the Pvr was severely impaired in the SNAP25 D51N and E55Q single mutants and D51N/E52Q double mutant (Fig. 1G). In contrast, the SNAP25 E52Q and the syntaxin-1 E228Q/D231N and D231N/E234Q/E238Q mutants displayed an enhanced Pvr (Fig. 1G,O), indicating a disinhibition phenotype. In analogy to the release probability, spontaneous release activity was increased for the SNAP25 E52Q mutant as well as for the syntaxin-1 E228Q/D231N double mutant and the D231N/E234Q/E238Q triple mutant, while the SNAP25 D51N or E55Q single mutants showed no change in spontaneous release (Fig. 1H,I,P,Q). Interestingly, the SNAP25 D51N/E52Q double mutant showed increased spontaneous release even though it showed severely impaired evoked release probability (Fig. 1G,H,I).

**Figure 1:**
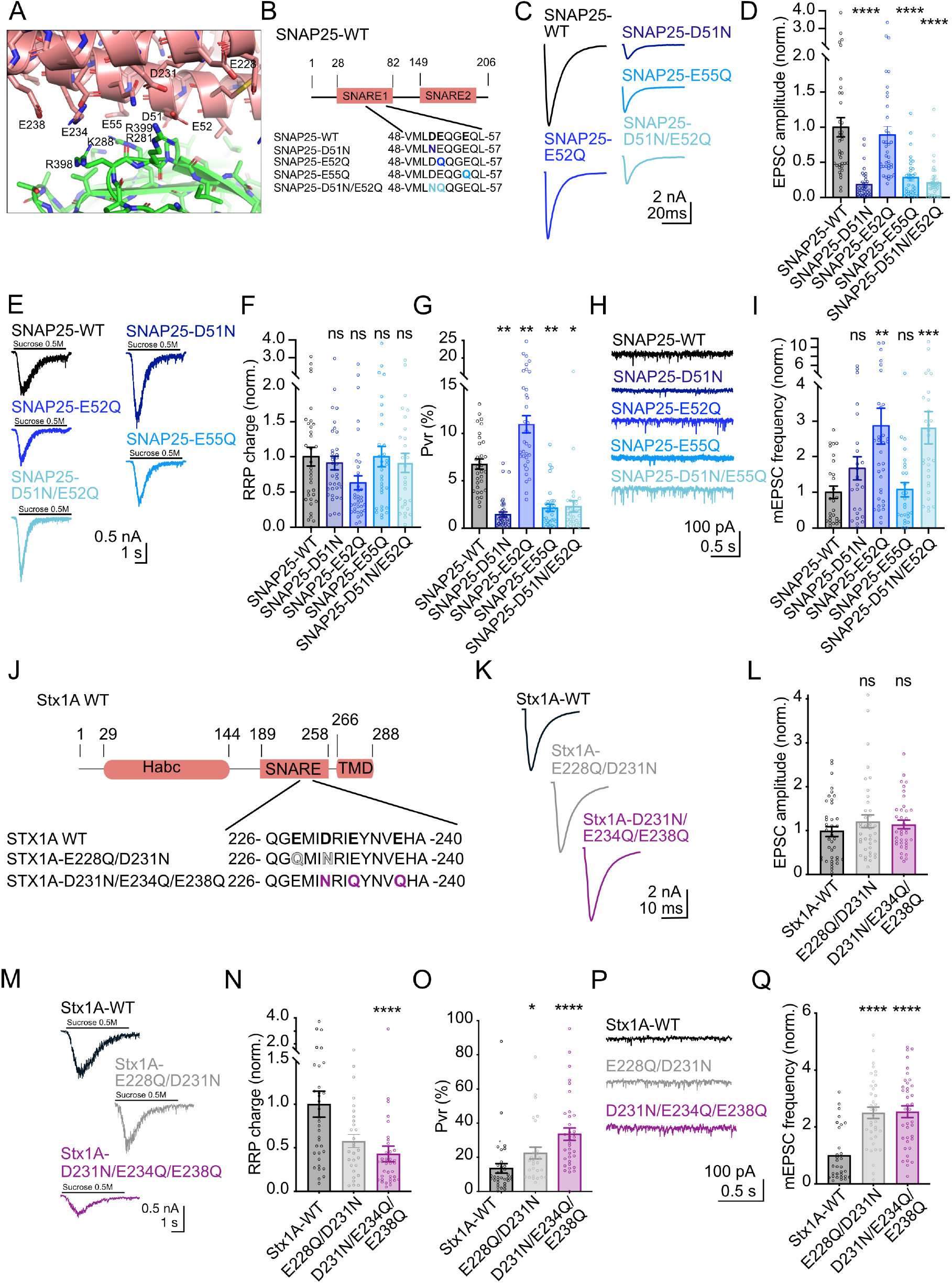
Syntaxin 1 and SNAP25 residues of the region II Syt1-SNARE complex interface mediate distinct pre-Ca^2+^ and post-Ca^2+^ functions. **(A),** Close-up view of region II of the primary interface. Proteins are represented by ribbon diagrams and stick models with nitrogen atoms in dark blue, oxygen in red, sulfur in yellow orange and carbon in salmon color (SNARE complex) or green (Syt1 C_2_B domain). Selected residues are labeled. (**B)** Schematics of SNAP25 mutations utilized. (**C)** Example traces and (**D)** quantification of the EPSC amplitude recorded for autaptic hippocampal neurons obtained from rescue experiments with SNAP25-WT, SNAP25-D51N, E52Q, E55Q, or D51N/E52Q. **(E)** Example traces and (**F)** quantification of the readily releasable pool (RRP) charge induced by 500 mM sucrose application obtained from the same neurons as in (C). (**G)** Quantification of the vesicle release probability (Pvr). (**H)** Example traces and (**I)** quantification of the miniature EPSC frequency (mEPSC) obtained from the same neurons as in (C). (**J)** Schematics of Stx1A mutations utilized. (**K)** Example traces and **(L)** quantification of the EPSC amplitude recorded for autaptic hippocampal neurons obtained from rescue experiments with Stx1A-WT, Stx1A-E228Q/D231N and Stx1A-D231N/E234Q/E238Q. **(M)** Example traces and **(N)** quantification of the readily releasable pool (RRP) induced by 500 mM sucrose application obtained from the same neurons as in (K). **(O)** quantification of the vesicle release probability (Pvr). **(P)** Example traces and **(Q)** quantification of the miniature EPSC frequency (mEPSC) obtained from the same neurons as in (K). Each data point represents a single recorded neuron. Between 35 and 39 neurons per group from 3 independent cultures were recorded and are shown as mean +/- SEM. Normalization in this and subsequent experiments was computed by dividing response from each neuron against mean values of the WT rescue group for each individual culture. ns: not significant, *p<0.05, **p<0.01, ***p<0.001 and ****p<0.0001.

Overall, these results further support the functional importance of the Syt1/SNARE complex primary interface. The differential phenotypes fit with previous observations that this interface contributes to functions before action potential triggering, i.e. SV priming and clamping of spontaneous release, or after action potential triggering, i.e. EPSC size and release probability. When mapping the phenotypes to the region II interface, it is striking that the two SNAP25 mutations that impair evoked release involve residues on one side of the SNAP25 N-terminal SNARE motif helix (D51N, E55; Fig 1G), whereas mutations that impair spontaneous fusion clamping and vesicle priming and boost release probability involve residues on the other side of this helix (SNAP25 E52, syntaxin-1 D231, E234, E238) (Fig. 1A,M,O,Q). These findings raise the intriguing hypothesis that the region II of the primary interface is composed of two functional interfaces, regulating either pre-Ca^2+^ or post-Ca^2+^ functions.

### The three Syt1 arginines in region II are important for Ca^2+^-triggered release

In the crystal structures of Syt1-SNARE complexes, SNAP25 E52 and syntaxin-1 D231, E234 and E238 are close to R398 and R399 of the Syt1 C_2_B domain, while SNAP25 D51 and E55 interact with other basic residues of Syt1, R281 and K288 (Fig. 1A). However, there is variability among these interactions in the various crystal structures that have been described (15, 19) and SNAP25 E55 interacts very often with R398 in MD simulations of the primed state (24)[Fig. S1A-C; see more detailed analysis in the accompanying paper (20)]. Hence, it is unclear to what extent mutations in the three arginines may also exhibit both pre- and post-Ca^2+^ dependent behavior. R398 and R399 mutants have been characterized using rescues in Syt1 KO neurons and *in vitro* liposome fusion assays, and were shown to be important for spontaneous fusion and for Ca^2+^-triggered release (13, 15, 19, 25) as well as for SV docking/priming (7). Since we previously did not examine the specific roles of those residues in spontaneous release, and previous experiments were performed in presence of Syt7, we repeated the R398Q, R399Q mutagenesis experiments in the Syt1/7 DKO background (Fig. 2). In addition, we also studied R281A and K288A single mutants as well as a R281A/K288A double mutant (Fig. 3).

**Figure 2:**
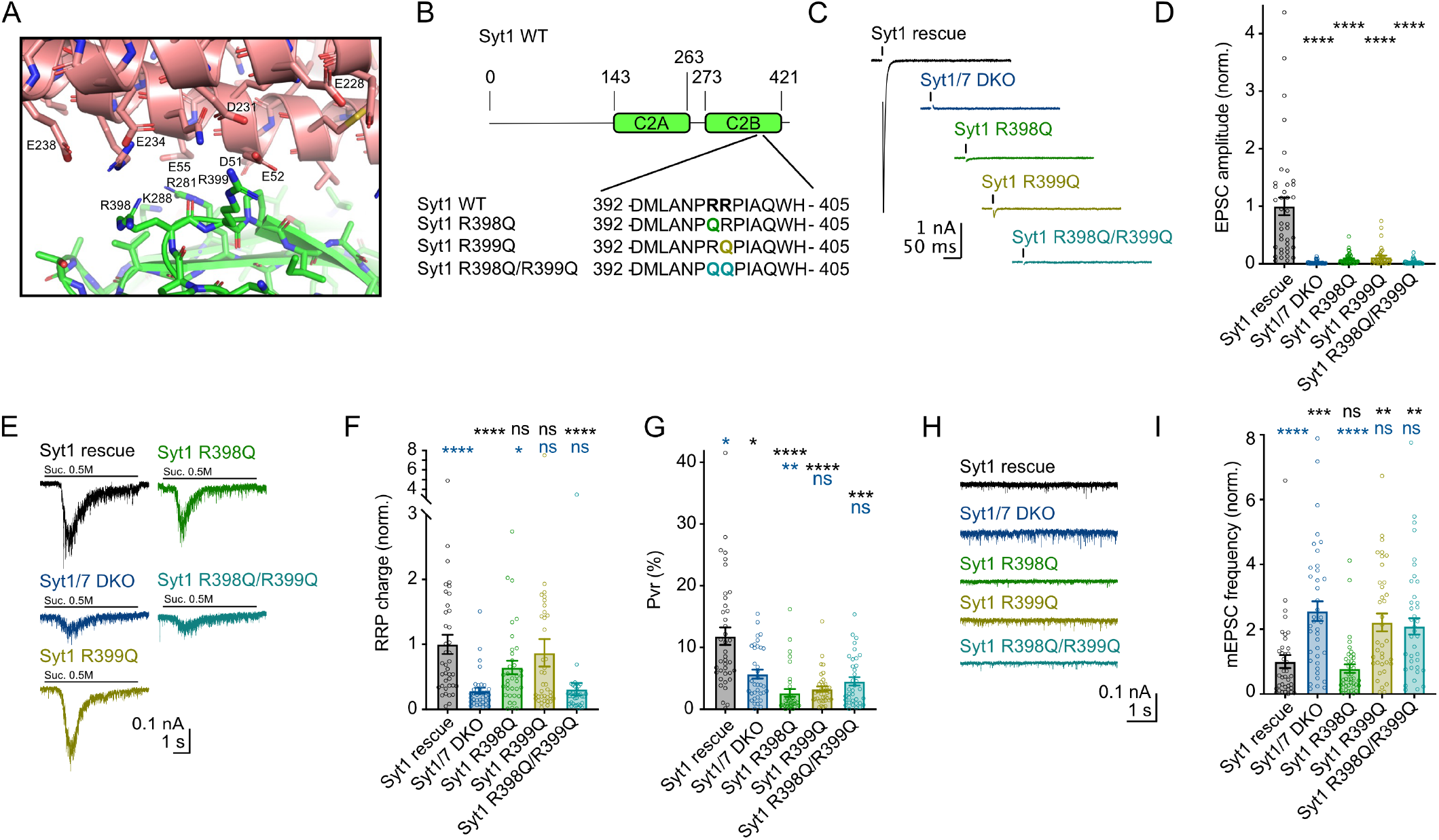
Region II Syt1 R398Q, R399Q mutants show distinct and overlapping roles in pre-Ca^2+^ and post-Ca^2+^ functions of Syt1. **(A)** Close-up view of Region II of the primary interface. Proteins are represented by ribbon diagrams and stick models with nitrogen atoms in dark blue, oxygen in red, sulfur in yellow orange and carbon in salmon color (SNARE complex) or green (Syt1 C_2_B domain). Selected residues are labeled. **(B)** Schematics of Synaptotagmin1 (Syt1) mutations utilized. **(C)** Example traces and **(D)** quantification of the EPSC amplitude recorded for Syt1/7 DKO autaptic hippocampal neurons rescued with Syt1 WT, Syt1 R398Q, Syt1 R399Q, or Syt1 R398Q/R399Q **(E)** Example traces and **(F)** quantification of the readily releasable pool (RRP) charge induced by 500 mM sucrose application obtained from the same neurons as in (C). **(G)** Quantification of the vesicle release probability (Pvr). **(H)** Example traces and **(I)** quantification of the miniature EPSC frequency obtained from the same neurons as in (C). Each data point represents a single recorded neuron. Between 39 and 42 neurons per group from 3 independent cultures were recorded and are shown as mean +/- SEM. ns: not significant, *p<0.05, **p<0.01, ***p<0.001 and ****p<0.0001.

**Figure 3:**
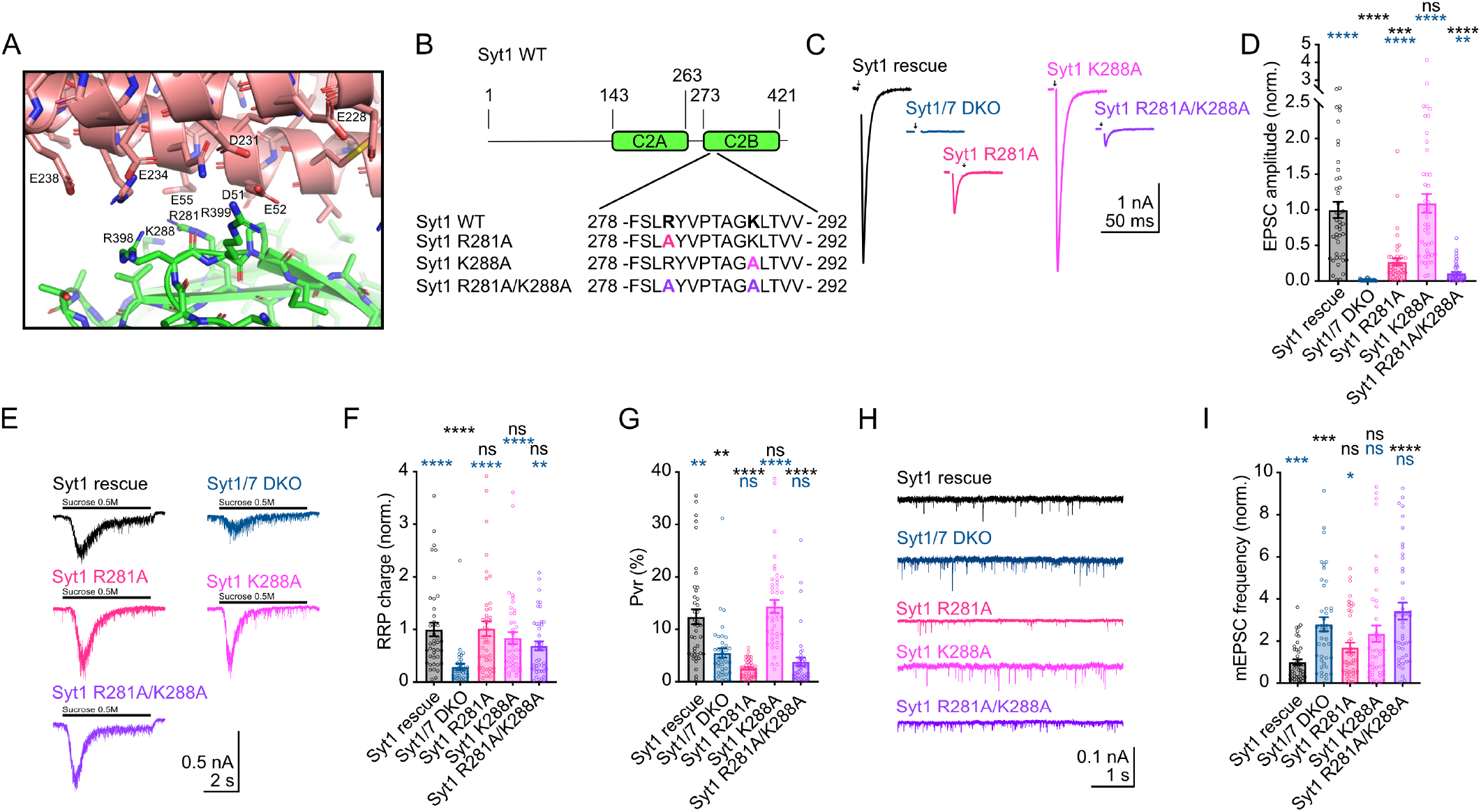
Region II Syt1 R281, R288 mutant analysis show selective loss of post-Ca^2+^ function of Syt1. **(A)** Close-up view of Region II of the primary interface. Proteins are represented by ribbon diagrams and stick models with nitrogen atoms in dark blue, oxygen in red, sulfur in yellow orange and carbon in salmon color (SNARE complex) or green (Syt1 C_2_B domain). Selected residues are labeled. **(B)** Schematics of Synaptotagmin1 (Syt1) mutations utilized. **(C)** Example traces and **(D)** quantification of the EPSC amplitude recorded for Syt1/7 DKO autaptic hippocampal neurons rescued with Syt1 WT, Syt1 R281A, Syt1 K288A, or Syt1 R281A/K288A. **(E)** Example traces and **(F)** quantification of the readily releasable pool (RRP) charge induced by 500 mM sucrose application obtained from the same neurons as in (C). **(G)** Quantification of the vesicle release probability (Pvr). **(H)** Example traces and **(I)** quantification of the miniature EPSC frequency obtained from the same neurons as in (C). Each data point represents a single recorded neuron. Between 44 and 45 neurons per group from 3 independent cultures were recorded and are shown as mean +/- SEM. ns: not significant, *p<0.05, **p<0.01, ***p<0.001 and ****p<0.0001.

Evoked responses were massively impaired in both R398Q, R399Q single mutants and the double R398Q/R399Q mutant, consistent with previous findings (13) (Fig. 2C,D,S3F). Analysis of RRP size showed that the R398Q/R399Q double mutant exhibited full loss of Syt1 priming function, consistent with previous work (7), while the Syt1 R398Q and R399Q single point mutants showed normal RRP size (Fig. 2E,F). Vesicular release probability was impaired in both single and the double point mutants (Fig. 2G), resulting in enhanced facilitation of EPSC amplitudes when five were evoked at a frequency of 50Hz (Fig. S3H). Analysis of spontaneous release showed that the double mutant R398Q/R399Q exhibited a 2.5-fold increased spontaneous release activity, while the R398Q mutant was WT-like and the R399Q mutant showed a similar increase in mEPSC frequency to that of the double mutant (Fig. 2H,I). Amplitudes of mEPSCs were not affected by the mutations (Fig. S3G). These results show that, together, R398/399 are essential for both pre-Ca^2+^ and post-Ca^2+^ functions. The phenotypes of individual point mutants are more discrete, as both mutants impair Ca^2+^- triggered release but not SV priming, and only the R399Q mutation impairs the clamping function.

Analysis of R281A, K288A single and double mutants showed that only R281 is critical for Ca^2+^-triggered release (Fig. 3C,D,S3C). Vesicle priming as measured by RRP size showed that single and double mutants had WT-like RRP sizes, although the double mutant had a tendency toward a reduced RRP size (Fig. 3E,F). Vesicular release probability was greatly impaired (Fig. 3G) supported by the analysis of short-term plasticity characteristics, which showed increased facilitation (Fig. S3E). Spontaneous release in the R281A and K288A single mutants were not statistically different from WT but showed a tendency to be higher, while the R281A/K288A double mutant showed a significant loss of clamping of spontaneous release (Fig. 3H,I). Again, mEPSC amplitudes were not affected in these mutants (Fig. S3D). Taken together, these results show that R281 is particularly important for the post-Ca^2+^ function of Syt1, but the R281A/K288A double mutant data suggest that these residues contribute to a small extent to pre-Ca^2+^ functions of Syt1. The limited contribution of K288 is in retrospect not surprising because MD simulations of the primed state show that the K288 side chain is seldom in contact with acidic residues of the SNARE complex (Fig. S1C); instead, this side chain has a tendency to bind to the flat bilayer that mimics the plasma membrane in the simulations (20) (Fig. S1C), which may account for the contribution of K228 to the priming/clamping phenotypes observed for the R281A/K288A double mutant. Overall, the analysis of region II of the primary interface from the side of Syt1 generally supports its role in priming, clamping and Ca^2+^-triggering, and in addition, reinforces the notion that different portions of the interface contribute differentially to the pre-Ca^2+^ and post-Ca^2+^ functions.

### Region I of the primary interface mediates SV priming but needs to dissociate for Ca^2+^- triggering of release

Given that region II of the primary interface has dual functions in release, we suggest that the interface may not be static, but rather can switch between alternative orientations that mediate pre-Ca^2+^ and post-Ca^2+^ functions. To further investigate the putative dynamic nature of the interface, we examined the functional consequences of a Syt1 Y338D mutation in region I and a A402T mutation in the small hydrophobic pocket between regions I and II (Fig. 4A,B), both of which suppressed the lethality caused by expression of Syt1 bearing C_2_B domain Ca^2+^- binding site mutations in *Drosophila* (21). We found that for both single point mutants and the double point mutant, the evoked response (Fig 4C,D,S4C) and vesicular release probability (Fig. 4G,S4E) were drastically impaired, although the A402T mutation displayed a slightly weaker phenotype than the other two mutants. Both pre-Ca^2+^ functions (RRP size and clamping of spontaneous release activity) were impaired in all these mutants (Fig. 4E,F,H,I), without affecting mEPSC amplitudes (Fig. S4D), demonstrating that disruption of region I or the hydrophobic pocket between regions I and II leads to a loss of pre-Ca^2+^ as well as post-Ca^2+^ functions.

**Figure 4:**
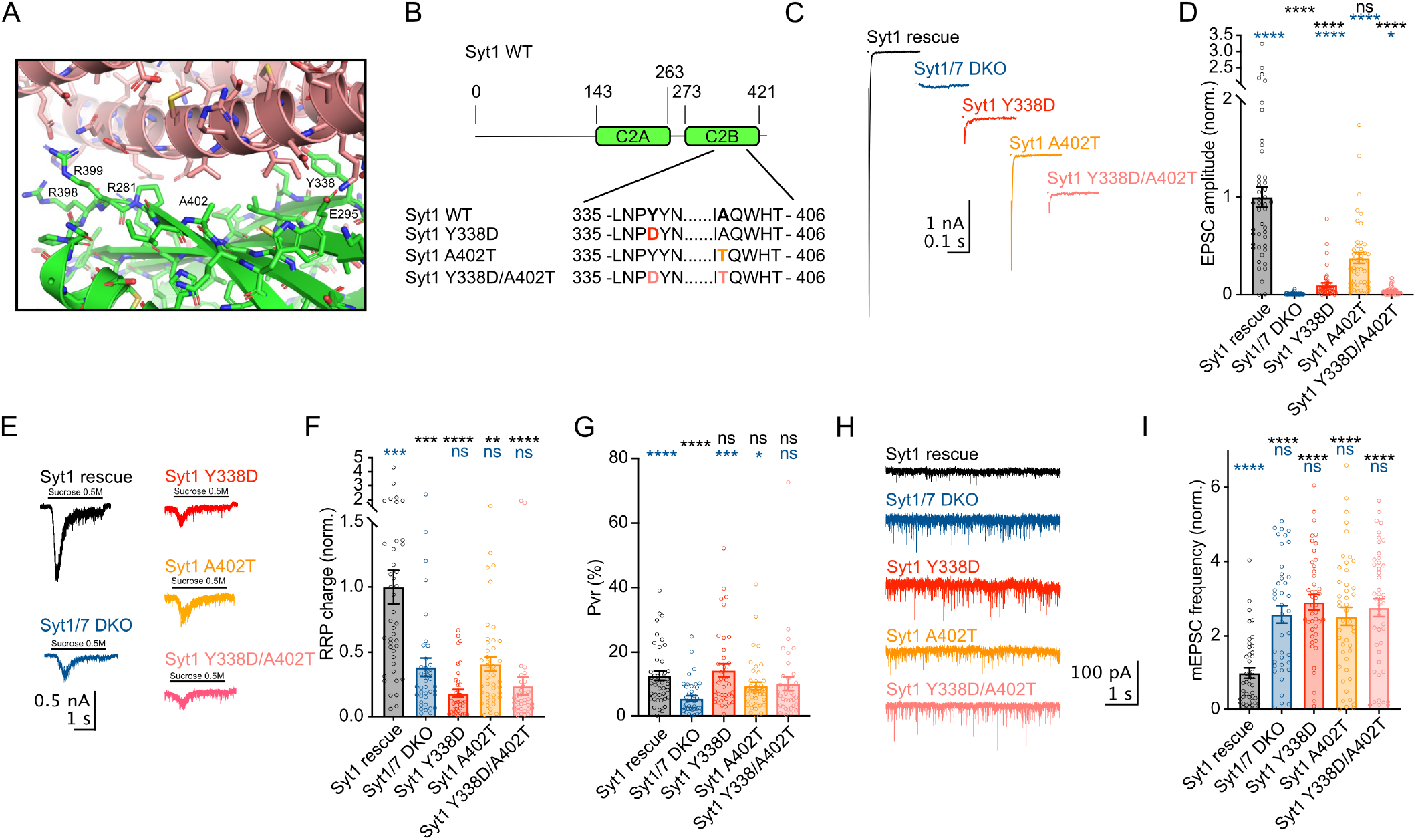
Region I Syt1 Y338D and A402T mutants display loss of pre-Ca^2+^ and post-Ca^2+^ functions of Syt1. **(A)** Close-up view of Region I and Region II of the primary interface. Proteins are represented by ribbon diagrams and stick models with nitrogen atoms in dark blue, oxygen in red, sulfur in yellow orange and carbon in salmon color (SNARE complex) or green (Syt1 C_2_B domain). Selected residues are labeled. **(B)** Schematics of Synaptotagmin1 (Syt1) mutations utilized. **(C)** Example traces and **(D)** quantification of the EPSC amplitude recorded for Syt1/7 DKO autaptic hippocampal neurons rescued with Syt1 WT, Syt1 Y338D, Syt1 A402T and Syt1 Y338D/A402T mutants. **(E)** Example traces and **(F)** quantification of the readily releasable pool (RRP) charge induced by 500 mM sucrose application obtained from the same neurons as in (C). **(G)** Quantification of the vesicle release probability (Pvr). **(H)** Example traces and **(I)** quantification of the miniature EPSC frequency obtained from the same neurons as in (C). Each data point represents a single recorded neuron. Between 41 and 45 neurons per group from 3 independent cultures were recorded and are shown as mean +/- SEM. ns: not significant, *p<0.05, **p<0.01, ***p<0.001 and ****p<0.0001.

Previous studies showed that an E295A/Y338W mutation in region I strongly impaired Ca^2+^- triggered release but preserved the ability of Syt1 to clamp spontaneous release (15) and, intriguingly, enhanced the affinity of the Syt1 C_2_B domain for the SNARE complex (17). To further understand the functional effects of this mutation and dissect the contributions of the two residue substitutions to these effects, we analyzed the phenotypes of the E295A/Y338W double mutant as well as E295A and Y338W single mutants (Fig. 5). The resulting phenotypes were strikingly different from the region I loss of function phenotypes (Fig. 4). While the evoked response for the E295A mutant was 90% reduced (Fig. 5C,D,S5C), the RRP charge was 50% larger than WT rescue (Fig 5E,F), and the clamping function was massively increased, as the spontaneous release rate was 4-fold lower than that of the Syt1 WT rescue (Fig. 5O). Upon closer inspection of mEPSC events, we found that the E295A mutant exhibited an approximately one-third reduction in mEPSC amplitude (Fig. 5N) and charge compared to WT rescue (Fig. S5D), indicating a possible impairment of postsynaptic glutamate receptor function (26, 27). Since mEPSC charge defines the contribution of individual vesicle fusion events to RRP estimates, the RRP size corrected by mEPSC charge was approximately twice the size in the E295A mutant compared to control (Fig. 5H). While the RRP size in the E295A mutant was increased, spontaneous release rates (Fig. 5O) and evoked release was decreased (Fig. 5G), arguing for a higher threshold for primed vesicles to transition to fusion. We tested for the decreased vesicle fusogenicity by the analysis of release kinetics induced by the hypertonic stimulus (28), which indeed was slowed down (Fig. 5 I-K), indicative of a reduced fusogenicity of primed vesicles. The phenotypes observed for the E295A/Y338W double mutant were very similar to those caused by the E295A mutation while no overt phenotypes were caused by the single Y338W mutation, showing that this substitution does not substantially alter Syt1 function (Fig. 5C-O). Interestingly, NMR experiments described in the accompanying paper showed that the E295A mutation enhances the affinity of the Syt1 C_2_B domain for the SNARE complex about 5- to 6-fold whereas the Y338W mutation has no effect (20). Overall, these results suggest that the E295A mutation causes a super primed, super clamped phenotype, essentially a gain of function phenotype for the pre-Ca^2+^ function of the primary interface. Given the observed lowered vesicle fusogenicity, we conclude that this mutant locks the primary interface in the pre-Ca^2+^ state, preventing the transition into the post-Ca^2+^ triggered state.

**Figure 5:**
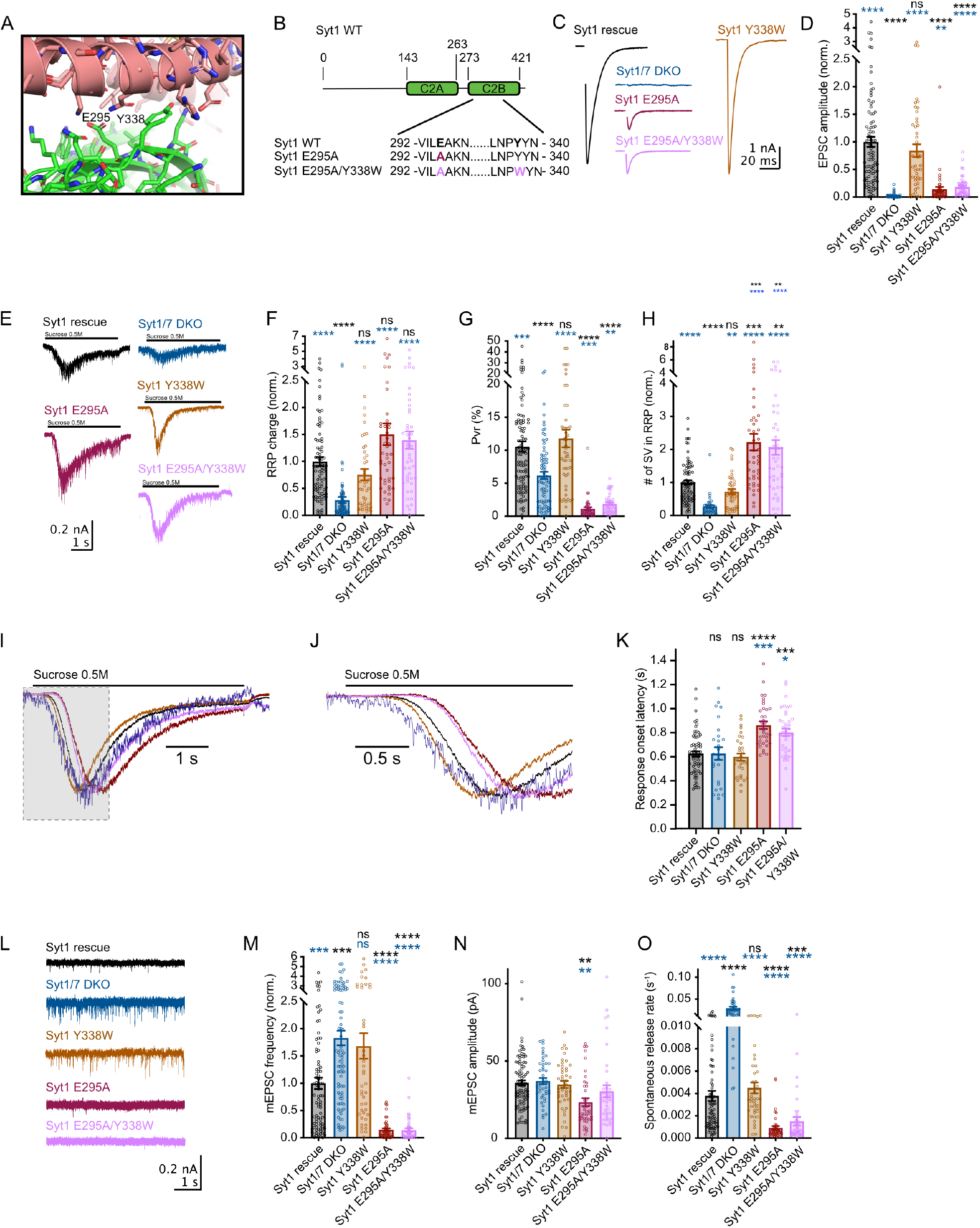
Region I Syt1 mutant E295A stabilizes the primed state and impairs Ca^2+^ triggered release. **(A)** Close-up view of Region I of the primary interface. Proteins are represented by ribbon diagrams and stick models with nitrogen atoms in dark blue, oxygen in red, sulfur in yellow orange and carbon in salmon color (SNARE complex) or green (Syt1 C_2_B domain). Selected residues are labeled. **(B)** Schematics of Synaptotagmin1 (Syt1) mutations utilized. **(C)** Example traces and **(D)** quantification of the EPSC amplitude recorded for Syt1/7 DKO autaptic hippocampal neurons rescued with Syt1 WT, Syt1 Y338W, Syt1 E295A, and Syt1 E295A/Y338W mutants **(E)** Example traces and **(F)** quantification of the readily releasable pool (RRP) charge induced by 500 mM sucrose application obtained from the same neurons as in (C). **(G)** Quantification of the vesicle release probability (Pvr). **(H)** Quantification of the number of synaptic vesicles (SV) in the RRP. **(I)** Average traces and **(J)** close-up view of synaptic responses induced by an application of 500 mM sucrose for Syt1 WT (n = 65), Syt1/7 DKO (n = 26), Syt1 Y338W (n = 32), Syt1 E295A (n = 35) and Syt1 E295A/Y338W (n = 38) mutants. **(K)** Quantification of the response onset latency normalized to Syt1 rescue. (**L)** Example traces and quantification of the miniature EPSC frequency **(M)**, amplitude **(N)** and the spontaneous release rate as a ratio of mEPSC frequency and the number of SV in the RRP **(O)** obtained from the same neurons as in (C). Each data point represents a single recorded neuron. Between 45 and 46 neurons per group from 3 independent cultures were recorded and are shown as mean +/- SEM. ns: not significant, *p<0.05, **p<0.01, ***p<0.001 and ****p<0.0001.

## Discussion

The synaptic SNARE proteins and Syt1 play essential roles in the process of exocytosis, and they do so in multiple steps from preparing SVs for fusion competence, to increasing signal to noise through suppression of spontaneous release, to ultimately mediating vesicle fusion at submillisecond speed in response to incoming action potentials. Previous studies showed that binding of Syt1 C2B domain to the SNARE complex via the primary interface is critical for neurotransmitter release and probably mediates more than one of these steps (13, 15, 19, 21), but the specific role in each step was unclear, particularly in Ca^2+^-triggering of release (29). In fact, some data suggested that Ca^2+^-binding to Syt1 leads to dissociation from the SNARE complex (17), consistent with the fact that the primary interface did not seem compatible with widespread models postulating that Syt1 facilitates membrane fusion by direct actions of its Ca^2+^-binding loops on the membranes [e.g. (30–33); see also (20)]. Our systematic analysis of primary interface mutants, together with correlations between our data and the effects of the same mutations on Syt1 C_2_B domain-SNARE complex binding (20), now consolidate the notion that the primary interface is critical for the functions of Syt1 in SV priming and in clamping of spontaneous release. Importantly, our results firmly establish that the primary interface is directly involved in Ca^2+^-triggering of neurotransmitter release and that this function involves a rearrangement of this interface in which region I dissociates but region II remains bound. Such a rearrangement is a central aspect of a model postulating that, upon Ca^2+^ binding, Syt1 pulls from the SNARE complex to facilitate jxt linker zippering and fast membrane fusion that we present in the accompanying paper (Fig. 6G; (20)).

**Figure 6.**
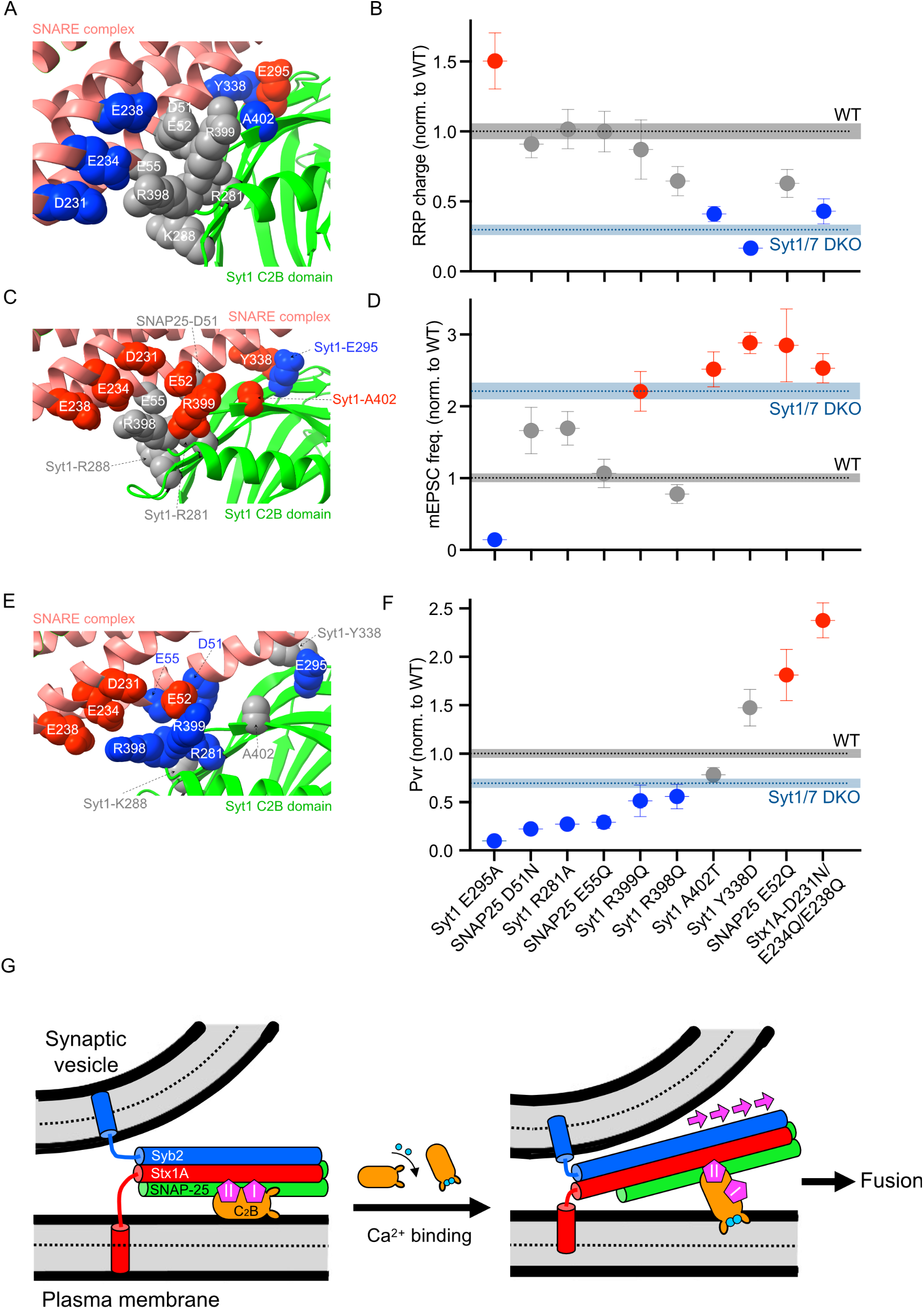
Summary of primary interface mutational phenotypes and a putative model of Syt1-mediated Ca^2+^-triggered SV fusion. **(A)** Map of residues that **(B)** modulate priming function. **(C)** Map of residues that **(D)** modulate spontaneous fusion clamping function. **(E)** Map of residues that modulate **(F)** Ca^2+^-triggered release probability. In **A, C** and **E,** molecular graphics of the C2B domain-SNARE complex interaction from PDB accession code 5CCH (15). Proteins are represented by ribbon diagrams and sphere models. Carbons are represented in salmon color (SNARE complex) or green (Syt1 C2B domain). Selected residues are labeled in red if they show a significantly increased response compared to WT rescue, grey if they are not significantly different from WT rescue and blue if they are significantly decreased compared to their respective WT rescue protein. In **B, D** and **F,** residues are labeled in red if they show a significantly increased response compared to WT rescue, grey if they are not significantly different from WT rescue and blue if they are significantly decreased compared to their respective WT rescue protein. The black dotted line represents WT rescue whereas the dark blue dotted line symbolizes the result for Syt1/Syt7 DKO response. **(G)** Model of how Syt1 triggers neurotransmitter release. In the primed, fusion clamped state of the release apparatus, the Syt1 C_2_B domain (orange) binds to the SNARE complex through the primary interface and to the plasma membrane through a polybasic region. Regions I and II are represented schematically with pink diamond shape forms. Binding of Ca^2+^ (blue circles) induces reorientation of the C_2_B domain to allow insertion of both Ca^2+^ binding loops into the plasma membrane and coordination of the Ca^2+^ ions by the C_2_B domain ligands and phospholipid head groups. Because of the reorientation, region I dissociates, but region II remains in contact, which in turn communicates a “rowing force” from the Syt C2B reorientation onto the SNARE complex that facilitates extension of the synaptobrevin and syntaxin-1 helices into the jxt linkers, which leads to fast membrane fusion.

The importance of the Syt1-SNARE complex primary interface for SV priming was supported by the strong impairments in the ability of Syt1 to rescue the RRP caused by a quintuple R281A/E295A/Y338W/R398A/R399A mutation (15) and by the R398Q/R399Q mutation (7) in this interface. However, the quintuple mutation combined residue substitutions that decrease (R281A, R398A, R399A) or increase (E295/Y338W) C_2_B-SNARE complex binding (17) and R398/R399 has been implicated in the ability of Syt1 to bridge membranes (30). Our results now show that rescue of the RRP is impaired not only by the R398Q/R399Q mutation in region II (Fig. 2E,F), which abolishes C_2_B-SNARE complex binding (17), but also by the Y338D and A402T mutations (Fig. 4E,F) that abrogate binding through region I but preserve binding through region II (20). Conversely, the E295A and E295A/Y338W mutations enhance binding through the primary interface ((17), (20) and both exhibit increased RRP size (Fig. 5E,F). The syntaxin-1 E228Q/D231N and D231N/E234Q/E238Q mutations in region II also impair priming (Fig. 1M,N). Although the syntaxin-1 mutations could also disrupt priming by weakening binding to Munc18-1 (34, 35), the overall data leave little doubt that the primary interface is crucial for the role of Syt1 in SV priming and show that both region I and region II are required for this function.

The phenotypes observed for the R281A/E295A/Y338W/R398A/R399A and R398Q/R399Q mutations (15) also suggested a key role for Syt1 in clamping spontaneous neurotransmitter release through the primary interface. This notion is also clearly strengthened by our data, including the enhancements in spontaneous release observed for many of the mutations in the primary interface that weaken or abolish Syt1-SNARE complex binding, and by the strong inhibition of spontaneous release caused by the E295A and E295A/Y338W mutations (Fig. 6C,D) that enhance the binding affinity (20). Analyses of the relationships between the parameters measured for the various mutants used in this study revealed a correlation between the ability of Syt1 mutants to restore the RRP (Fig. 6A,B) and to inhibit spontaneous release (Fig. 6C,D), the two pre-Ca^2+^ functions. The correlation is not perfect, but this is not surprising because enhancement of spontaneous release may involve more than one mechanism (see below).

Whereas both region I and region II are important for the pre-Ca^2+^ functions of Syt1, our data clearly establish that the post-Ca^2+^ function requires binding *via* region II and dissociation of region I, thus showing that a rearrangement of the primary interface must occur for Ca^2+^- triggering of neurotransmitter release. Particularly compelling evidence for the notion that region I must dissociate is the observation that the E295A and E295A/Y338W mutations that enhance C_2_B-SNARE complex binding (20) strongly impair Ca^2+^-evoked release even though they enhance priming (Fig. 5C-F; Fig 6E,F). The key role of region II for Ca^2+^-triggering of release is clearly shown by the specific post-Ca^2+^ effects of the R281A and R398Q in Syt1 C_2_B and the D51N and E55Q mutations in SNAP25, which do not affect SV priming or spontaneous release but strongly disrupt evoked release, severely decreasing the vesicular release probability (Fig. 1C-I, 2C-I, 3C-H; Fig. 6F). Conversely, the E52Q mutation in SNAP25 as well as the E228Q/D231N and D231N/E234Q/E238Q mutations in syntaxin-1 do not alter Ca^2+^-evoked release but enhance spontaneous release and/or impair vesicle priming (Fig. 6B,D). Moreover, they also enhance release probability, thus playing a disinhibitory role on release (Fig. 6F).

The locations of SNARE acidic residues in the states most often visited during MD simulations of primed Syt1-SNARE complexes bridging a vesicle and a planar bilayer ((24), (20) (see example in Fig. S1C) exhibit a striking pattern: R281 binds simultaneously to D51 and E55 of SNAP25 while R398 binds to E55 and other acidic residues of SNAP25 (D58 and E62) on one side of the N-terminal SNARE motif helix, which can be viewed as the activating face of region II (Fig. S1B,C); in contrast, E52 of SNAP25 and the syntaxin-1 acidic residues are located on the other side of this helix, which thus appears to act as an inhibitory face of region II (Fig. S1B,C). Consistent with this view, the D51N/E52Q mutation in SNAP25 yields mixed phenotypes, impairing evoked release but enhancing spontaneous release (Fig. 1C,D,H,I), because it involves residues from both faces. These results argue that structural intermediates can exist that are neither clamped nor efficiently Ca^2+^-triggered. The single R399Q mutation also yielded mixed phenotypes, as it severely disrupted evoked release but enhanced spontaneous release (Fig. 2C,D,H,I,S3F). Thus, R399 contributes to both activating and inhibitory interactions. Note also that R281 and R398 have not only post-Ca^2+^ but also pre-Ca^2+^ functions, as the R281A mutation enhances spontaneous release when combined with K288A (Fig. 3H,I), and R398Q impairs SV priming when combined with R399Q (Fig. 2E,F). The different functions may involve distinct configurations of the arginine-acidic residue interactions in region II. Indeed, diverse configurations of these interactions were observed in the various crystal structures of the SNARE complex that are available (15, 19) and during the MD simulations (Fig. S1A-C) (see (20) for a more detailed analysis), and re-modeling of these interactions during the rearrangements that take place upon Ca^2+^ influx should be facilitated by the abundance of SNARE acidic residues near the three arginines. Based on the observed phenotypes, it seems likely that configurations in which the arginines interact primarily with the activating face of region II (e.g. Fig. S1C) trigger neurotransmitter release upon Ca^2+^ influx, whereas configurations involving more interactions with the inhibitory face hinder Ca^2+^-evoked release.

How do these new insights help us to understand the overall changes of the release apparatus when transitioning from the primed/clamped to the Ca^2+^-activated state? We know that action potential induced Ca^2+^ influx triggers chelation of Ca^2+^ between the top loops of the C_2_B domain and the phospholipid membrane, and the insertion of both Ca^2+^-binding loops into the membrane (30, 36, 37), leading to an approximately perpendicular orientation with respect to the bilayer (38). Since binding of the C_2_B domain to the SNARE complex via the primary interface and to the membrane results in a parallel orientation (24), Ca^2+^ binding is expected to induce a reorientation of the C_2_B domain (Fig. 6G). We further know from the phenotypes of the E295A mutant that region I of the primary interface must dissociate to enable Ca^2+^ triggering of release, while region II stays in contact with the SNARE complex through the malleable interactions of the C_2_B arginines with the SNARE acidic residues. Modeling shows that, if C_2_B reorients but never loses contact with the SNARE complex, the reorientation may create a force that pulls the SNARE complex away from the center of the membrane-membrane interface where fusion should occur, leading in turn to a pulling force on the jxt linkers (Fig. 6G). This action may facilitate zippering of the linkers, which is critical for neurotransmitter release (3, 5), brings the TM regions to the membrane interface (39) and, based our recent MD simulations (6), can lead to fast, microsecond scale membrane fusion. The role of the Syt1 C_2_A domain in this model is unclear, but it is plausible that this domain also helps to pull the SNARE complex and/or provides a support point on the membrane to facilitate the application of force by the C_2_B domain [see a more detailed description of this lever model of Syt1 action in the accompanying paper (20)].

The finding that linker zippering is not crucial for spontaneous release (3, 5) suggests that there are important differences between the mechanisms of spontaneous and evoked release. It is plausible that spontaneous release requires dissociation of Syt1 from the SNAREs because it involves natural motions of the SNARE complex that lead to membrane fusion without linker zippering and that are prevented by simultaneous binding of Syt1 to the SNARE complex and the plasma membrane. This model can explain why spontaneous release is comparable to that observed for the Syt1/7 DKO for mutations that abolish the priming function of Syt1 and C_2_B-SNARE complex binding, such as R398Q/R399Q (17) (Fig. 2E-I). The model also accounts for the enhancement of spontaneous release caused by various mutations that interfere with the primary interface (Fig. 2H,I, 3H,I, 4H,I) and the inhibition of spontaneous release caused by the E295A and E295/Y338W mutations that enhance C_2_B-SNARE complex binding (Fig. 5L-O). However, the R281A and R398Q mutations do not enhance spontaneous release (Fig. 2H,I, 3H,I, Fig 6D) even though they impair C_2_B-SNARE complex binding ((17) (20). Since R281 and R398 are key for the C_2_B reorientation that leads to Ca^2+^-evoked release, it is plausible that spontaneous release may occur by more than one mechanism, one involving Ca^2+^-independent reorientation of C_2_B and the other through dissociation from the SNARE complex, and that the R281A and R398Q mutations hinder the former but facilitate the latter, leading to cancellation of opposing effects. The notion that similar mechanisms can induce spontaneous and evoked release is supported by the finding that both forms of release are similarly enhanced by tryptophan mutations in the Ca^2+^-binding loops (11) and can also explain the increase in spontaneous release caused by a mutation in the aspartate Ca^2+^-ligands of the Syt1 C_2_B domain that abolishes Ca^2+^-binding to C_2_B and Ca^2+^-evoked release (19), as neutralization of the aspartates should favor Ca^2+^-independent C_2_B reorientation by decreasing electrostatic repulsion of the Ca^2+^-binding region with the lipids (36). It is also possible that spontaneous release in the Syt1/7 DKO or the R398Q/R399Q mutant that cannot bind to the SNARE complex involves also more than one mechanism mediated by primed SNARE complexes that are or are not bound to another Ca^2+^ sensor.

In summary, multiple aspects of the molecular mechanisms underlying Syt1 action remained to be fully understood, but extensive experimental data available on Syt1 and the SNAREs, together with the convergence of the electrophysiological data presented here and the biophysical studies described in the accompanying paper, provide compelling evidence for a model in which the rearrangement of the primary interface between Syt1 and the SNARE complex is a key event for neurotransmitter release. Specifically, we suggest that Syt1 and the SNARE complex maintain contact while Ca^2+^ triggers reorientation of Syt1 on the membrane. The force resulting from the parallel-to perpendicular reorientation of the Syt1 C2B domain is transferred to the SNARE complex that in turn exerts a pulling force onto the jxt linker and the transmembrane domains of the SNARE proteins, initiating vesicle fusion.

## Materials and Methods

### Animal maintenance and mouse lines

All mouse experiments were performed in accordance with the regulation of the animal welfare committee of the Charité – Universitätsmedizin Berlin. Time pregnant females were anesthetized and euthanized at E18 as permitted by the Landesamt für Gesundheit und Soziales (LaGeSo) Berlin under the license number T0220/09 and G106/20.

### Primary hippocampal cultures

Primary murine hippocampal neurons were prepared from newborn (P0-P2) for Stx1A/STX1B DKO (40) and cSyt1/Syt7 DKO mice (24) (gift from Prof. Thomas Südhof, Stanford University) or E18 embryonic mice for SNAP25 KO mice (41) of either sex, as described previously (42). Briefly, hippocampi were dissected, and neurons dissociated by an enzymatic treatment using 25 units per ml of papain for 45 min at 37 °C. For electrophysiology, low-density cultures of 3 x 10^3^ neurons/well were seeded on astrocyte micro-islands (35 mm diameter) for autaptic cultures. Astrocyte feeder layers were prepared 1-2 weeks before neuronal seeding. For Western blots (WB), hippocampal neurons were plated onto a continental astrocyte feeder layer at a density of 100 x 10^3^ neurons/well in a 6 well plate. After plating, neurons were incubated in Neurobasal-A medium (Invitrogen) supplemented with B-27 (Invitrogen), 50 μg/ml streptomycin and 50 IU/ml penicillin at 37 °C. Electrophysiological, imaging and Western blot experiments were performed at DIV 13-20.

### Lentiviral constructs and virus production

For expression of syntaxin-1, SNAP25 or Syt1 variants within neuronal cells, modified lentiviral vectors were used. All lentiviral constructs were generated through the Gibson assembly method (NEB) with the corresponding cDNAs and with a human synapsin-1 promoter-driven lentiviral shuttle vector (f(syn), based on FUGW (43)) that could contain either nuclear-localized (NLS) GFP or RFP that was fused C-terminally to a self-cleaving P2A peptide (44) to allow polycistronic translation.

Lentiviral particles were prepared by the Charité Viral Core Facility as previously described (43). Briefly, HEK293T cells were cotransfected with the shuttle vector f(syn)NLS-GFP-P2A-GOI-WPRE and helper plasmids, pCMVdR8.9 and pVSV.G with polyethylenimine. Virus containing supernatant was collected after 72 h, filtered, aliquoted, flash-frozen with liquid nitrogen, and stored at -80°C. For infection, about 5 x 10^5^ -1 x 10^6^ infectious virus units were pipetted onto 1 DIV hippocampal Stx1A/Stx1B DKO, SNAP25, or cSyt1/Syt7 DKO neurons per 35 mm-diameter well. In the case of Stx1A/STX1B DKO and cSyt1/Syt7 DKO hippocampal neurons Cre expressing virus was also added to the medium at DIV1.

### Electrophysiology

Whole-cell patch-clamp recordings were performed on autaptic cultures at room temperature at days in vitro 13-20 (42). Synaptic currents were recorded using a Multiclamp 700B amplifier (Axon Instruments) controlled by Clampex 9 software (Molecular Devices). A fast perfusion system (SF-77B; Warner Instruments) continuously perfused the neurons (1 - 2 ml/min) with an extracellular solution that contains the following (in mM): 140 NaCl, 2.4 KCl, 10 HEPES (Merck), 10 glucose (Carl Roth), 2 CaCl_2_ (Sigma-Aldrich), and 4 MgCl_2_ (Carl Roth) (∼300mOsm; pH 7.4). Somatic whole cell recordings were obtained using borosilicate glass pipettes, with a tip resistance of 2 - 5 MΩ and filled with an intracellular solution that contains the following (in mM): 136 KCl, 17.8 HEPES, 1 EGTA, 4.6 MgCl_2_, 4 Na_2_ATP, 0.3 Na2GTP, 12 creatine phosphate, and 50 U/ml phosphocreatine kinase (∼300 mOsm; pH7.4). Membrane capacitance and series resistance were compensated at 70% and only neurons with a series resistance lower than 12 MΩ were recorded further. Data were filtered by a low-pass Bessel filter at 3 kHz and sampled at 10 kHz using an Axon Digidata 1322A digitizer (Molecular Devices).

Neurons were clamped at -70 mV, and action potentials triggered by a 2 ms depolarization to 0 mV to measure EPSCs (excitatory postsynaptic currents). To quantify spontaneous release, 40 s of 1 kHz low pass filtered recordings in control and in glutamate receptor antagonist containing solutions were analyzed for the presence of mEPSC events using template algorithm based software in AxoGraph X (AxoGraph Scientific). mEPSCs were defined as events with 0.15 – 1.5 ms rise time and 0.5 – 5 ms half-width. False positive mEPSC events obtained in NBQX were subtracted to calculate the frequency of spontaneous events. The readily-releasable pool (RRP) was determined by applying hypertonic extracellular solution (included additional 500 mM sucrose) for 5 s and integrating the transient inward response component (23). The Pvr of each cell was calculated by dividing the average charge of the EPSC by the RRP charge. Spontaneous release rate was calculated by dividing the mEPSC frequency by the number of synaptic vesicles in the RRP. The number of synaptic vesicles in the RRP was calculated by dividing the RRP size by the mean mEPSC charge. To measure vesicle fusogenicity, we measured the response onset latency between the open tip control for solution exchange and the onset of the sucrose response (28).

### Western Blot

For detection of syntaxin1-A, SNAP25 or Syt1 protein levels by Western blotting, protein lysates were obtained from mass culture hippocampal neurons (DIV 13-16) grown on WT astrocyte feeder layers. Briefly, cells were lysed using 50 mM Tris/HCl (pH 7.9), 150 mM NaCl, 5 mM EDTA, 1 % Triton-X-100, 1 % Nonidet P-40, 1 % sodium deoxycholate, and protease inhibitors (complete protease inhibitor cocktail tablet, Roche Diagnostics GmbH). Proteins were separated by SDS-PAGE and transferred overnight at 4°C to nitrocellulose membranes. After blocking with 5 % milk powder (Carl Roth GmbH) for 1 hour at room temperature, membranes were incubated with mouse monoclonal anti-syntaxin-1A (1:10,000; Synaptic Systems), mouse anti-Syt1 (1:1,000; Synaptic Systems), mouse anti-SNAP25 (1:10,000; Synaptic Systems) and mouse monoclonal anti-betaTubulinIII (1:10,000; Sigma) antibodies for 1 hour at room temperature. The membranes were washed several times with PBS-Tween before being incubated with the corresponding horseradish peroxidase-conjugated goat secondary antibodies (all from Jackson ImmunoResearch Laboratories). Protein expression levels were visualized with ECL Plus Western Blotting Detection Reagents (GE Healthcare Biosciences).

### Structural representation of the primary interface

Molecular graphics of the SNARE complex and the C2B domain of Syt1 in Figure 6 were modelized and visualized with UCSF ChimeraX 1.7 (Pettersen et al., 2021 PMID: 32881101) using the coordinates of the atomic model corresponding to the accession code 5CCH deposited in the Protein Data Bank (15).

### Statistical Analysis

Statistical tests were performed with Prism 7 (GraphPad Software). For bar plots data are represented as mean ± SEM. First, all data were tested for normality using the D’Agostino & Pearson test. If they pass the parametric assumption, the Kruskal-Wallis test is performed followed by Dunn’s test. Data were considered statistically significant if p<0.05 and represented as follow: *p<0.05, **p<0.01, ***p<0.001 and ****p<0.0001. All experiments in the manuscript are performed on at least three cultures unless explicitly notified.

### Data and materials availability

Data and Materials of this study are available from the corresponding authors upon reasonable request.

## Acknowledgements

We thank Prof. Thomas Südhof for providing us with the cSyt1/Syt7 DKO mice. We are grateful to Berit Söhl-Kielczynski, Bettina Brokowski, Katja Pötschke and Heike Lerch for excellent technical assistance. We are thankful for the services of the Charité viral core facility for virus production. We thank Dr. Melissa Herman for critical reading and editing of this manuscript and the Rosenmund lab members for stimulating discussions. Molecular graphics were performed with UCSF ChimeraX, developed by the Resource for Biocomputing, Visualization, and Informatics at the University of California, San Francisco, with support from the National Institutes of Health R01-GM129325 and the Office of Cyber Infrastructure and Computational Biology, National Institute of Allergy and Infectious Diseases. This work was supported by the Deutsche Forschungsgemeinschaft projects 278001972, 184695641, EXC-2049-390688087; Ro1296 20-1 (all to CR) and NIH Research Project Award R35 NS097333 (to JR).

## Declaration of interests

The authors declare no competing interests.

## Notes

### Competing Interest Statement

The authors have declared no competing interest.

