## Supplementary materials for "Neurotransmitter release is triggered by a calcium-induced rearrangement in the Synaptotagmin-1/SNARE complex primary interface"

PNAS

**This PDF file includes:**

Figures S1 to S5

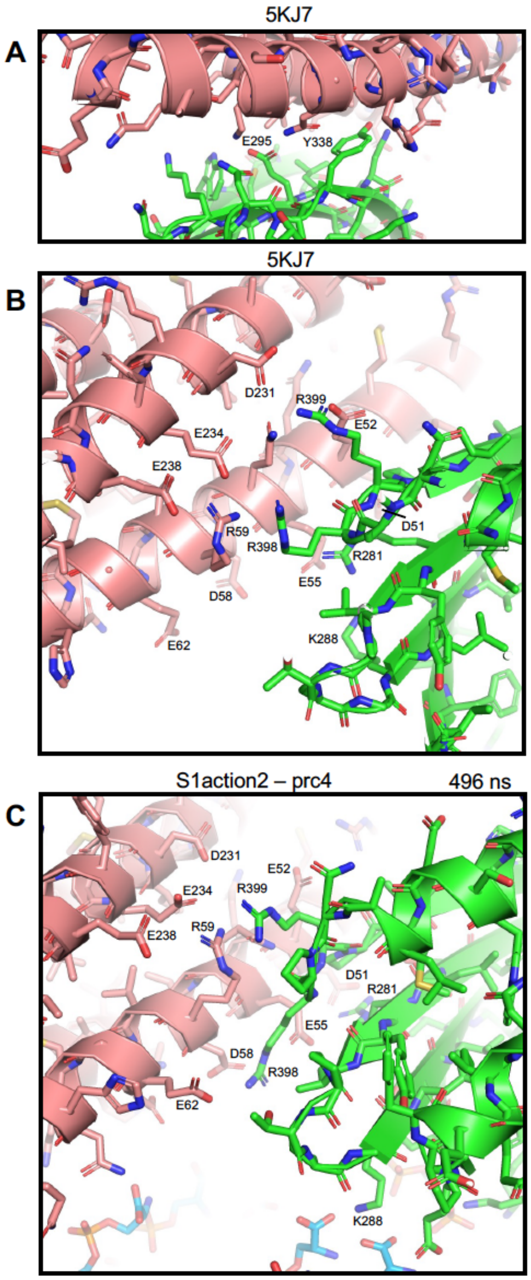


**Figure S1. Variability in region II of the primary interface.** Close-up views of region I (**A**) and region II (**B**) of the primary interface in a crystal structure of a Syt1-SNARE complex (PDB accession code 5KJ7). (**C**) Close-up view of one of the primed complexes (prc4) at 496 ns of the s1action2 simulation of the accompanying paper (1). Proteins are represented by ribbon diagrams and stick models with nitrogen atoms in dark blue, oxygen in red, sulfur in yellow orange and carbon in salmon color (SNARE complex) or green (Syt1 $C_2B$ domain). In (C), lipids from the flat bilayer are represented by stick models with carbon atoms in light blue, nitrogen atoms in dark blue, oxygen in red and phosphorous in orange. Acidic residues from the SNAREs as well as K288 and the three arginines of the $C_2B$ domain are labeled. Note that in (C) K288 of the $C_2B$ domain interacts with phospholipid head groups rather than the SNARE complex.

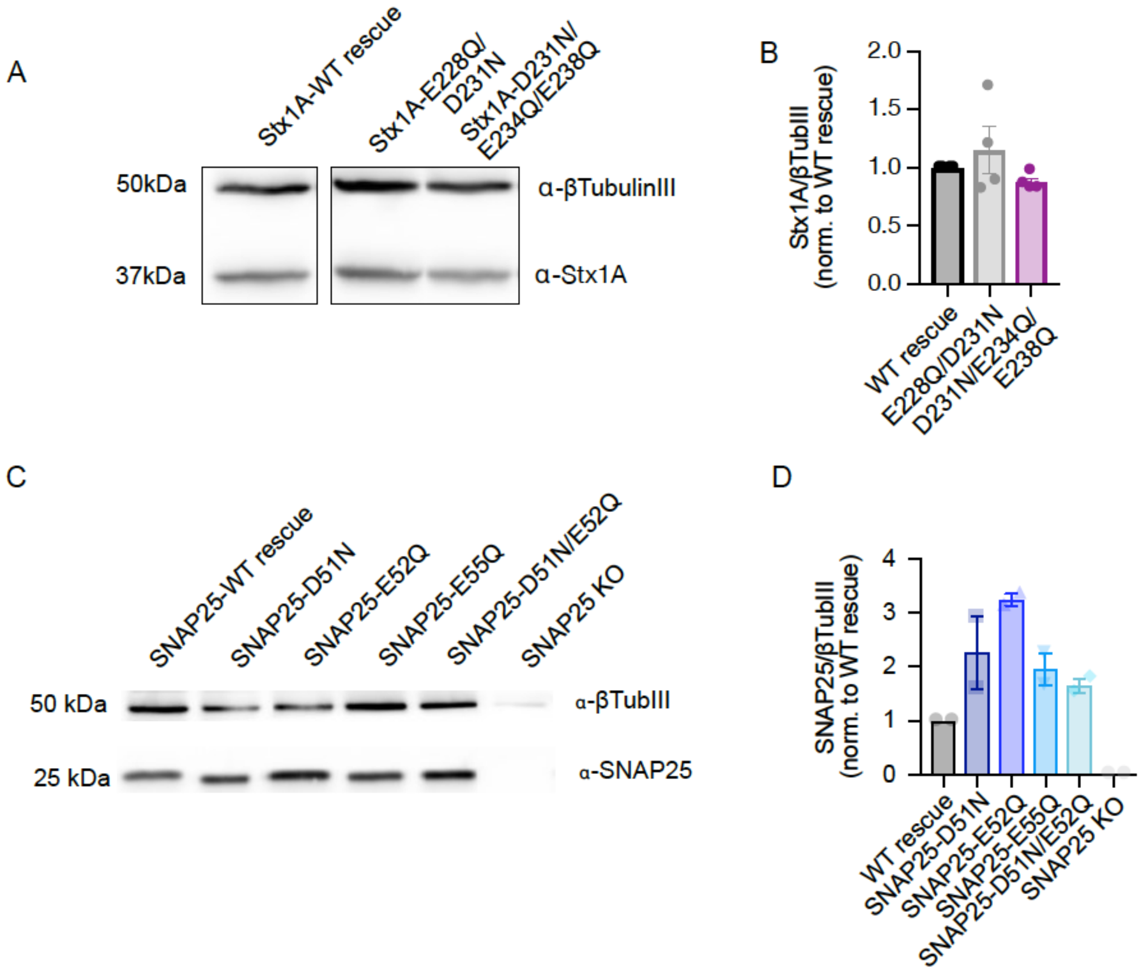


**Figure S2: Quantification of Stx1A and SNAP25 mutant levels i n hippocampal neurons.** (**A**) Example immunoblot and (**B**) quantification of protein levels for each experimental group. Proteins were detected using antibodies against beta-TubulinIII as loading control and Stx1A as the protein of interest. (**C**) Example immunoblot and (**D**) quantification of protein levels for each experimental group. Proteins were detected using antibodies against beta-TubulinIII as loading control and SNAP25 as the protein of interest. Protein expression for each mutant is normalized to their respective WT-rescue protein level. N = 2 – 4 replicates.

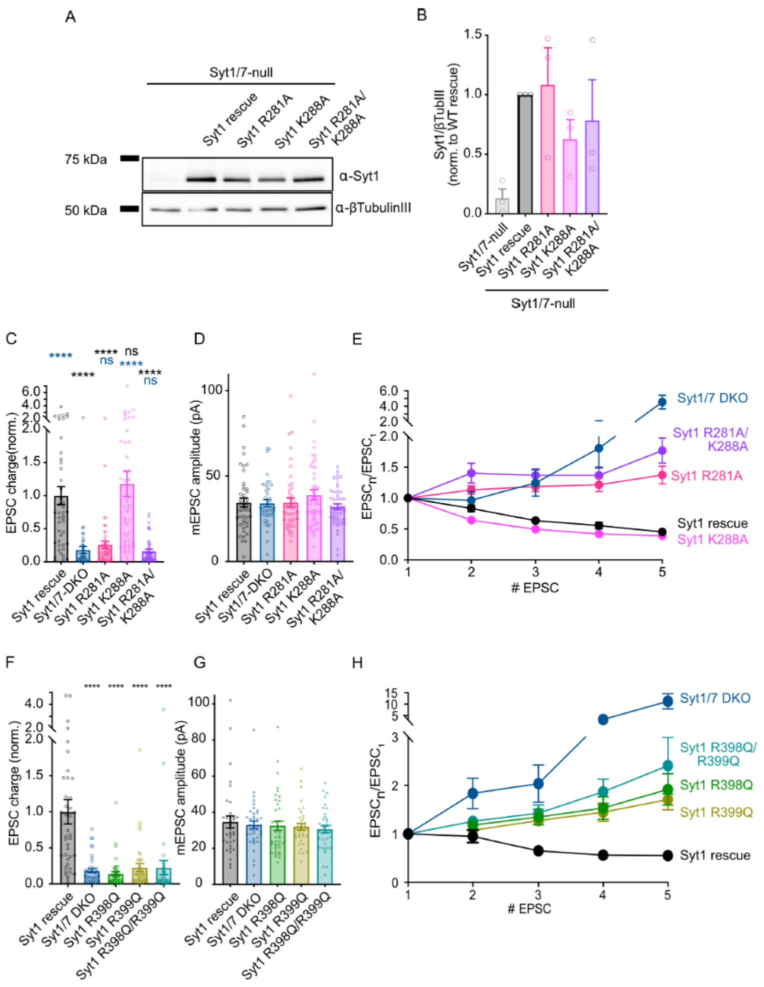


**Figure S3: Quantification of Syt1 mutant levels in hippocampal neurons and electrophysiological parameters of autaptic hippocampal mouse neurons rescued with Syt1 Region II mutants. (A)** Example immunoblot and **(B)** quantification of protein levels for each experimental group. Proteins were detected using antibodies against beta-TubulinIII as loading control and Syt1 as the protein of interest. Protein expression for each mutant is normalized to Syt1 WT-rescue protein level. n = 3 replicates. **(C)** Quantification of the EPSC charge recorded for Syt1/7 DKO autaptic hippocampal neurons rescued with Syt1 WT, Syt1 R281A, Syt1 K288A, or Syt1 R281A/K288A obtained from the same neurons as in (Fig. 3C). **(D)** Quantification of the miniature EPSC amplitude obtained from the same neurons as in (Fig. 3C). **(E)** Quantification of short-term plasticity measured by 5 stimulations at 50 Hz obtained from the same neurons as in (Fig. 3C). **(F)** Quantification of the EPSC charge recorded for Syt1/7 DKO autaptic hippocampal neurons rescued with Syt1 WT, Syt1 R398Q, Syt1 R399Q, or Syt1 R398Q/R399Q obtained from the same neurons as in (Fig. 2C). **(G)** Quantification of the miniature EPSC amplitude obtained from the same neurons as in (Fig. 2C). **(H)** Quantification of short-term plasticity measured by 5 stimulations at 50 Hz obtained from the same neurons as in (Fig. 2C). Between 39 and 45 neurons per group from 3 independent cultures were recorded and are shown as mean +/- SEM. ns: not significant, *p<0.05, **p<0.01, ***p<0.001 and ****p<0.0001.

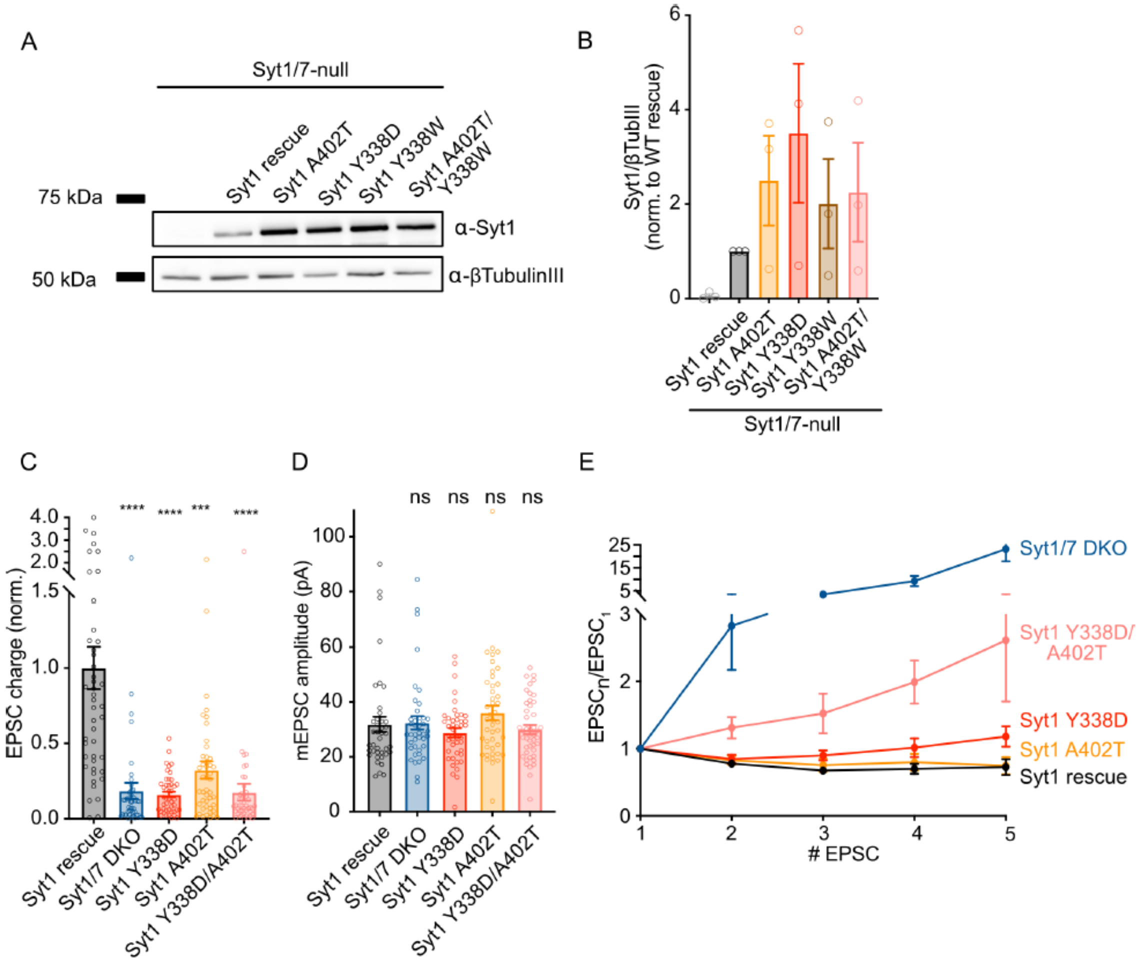


**Figure S4: Quantification of Syt1 mutant levels in hippocampal neurons and electrophysiological parameters of autaptic hippocampal mouse neurons rescued with Syt1 Y338D, Y338W, Syt1 A402T and Syt1 Y338D/A402T mutants from Region I.** (**A**) Example immunoblot and (**B**) quantification of protein levels for each experimental group. Proteins were detected using antibodies against beta-TubulinIII as loading control and Syt1 as the protein of interest. Protein expression for each mutant is normalized to Syt1 WT-rescue protein level. n = 3 replicates. (**C**) Quantification of the EPSC charge recorded for Syt1/7 DKO autaptic hippocampal neurons rescued with Syt1 WT, Syt1 Y338D, Syt1 A402T, or Syt1 Y338D/A402T obtained from the same neurons as in (Fig. 4C). **(D)** Quantification of the miniature EPSC amplitude obtained from the same neurons as in (Fig. 4C). **(E)** Quantification of short-term plasticity measured by 5 stimulations at 50 Hz obtained from the same neurons as in (Fig. 4C). Between 41 and 45 neurons per group from 3 independent cultures were recorded and are shown as mean +/- SEM. ns: not significant, *p<0.05, **p<0.01, ***p<0.001 and ****p<0.0001.

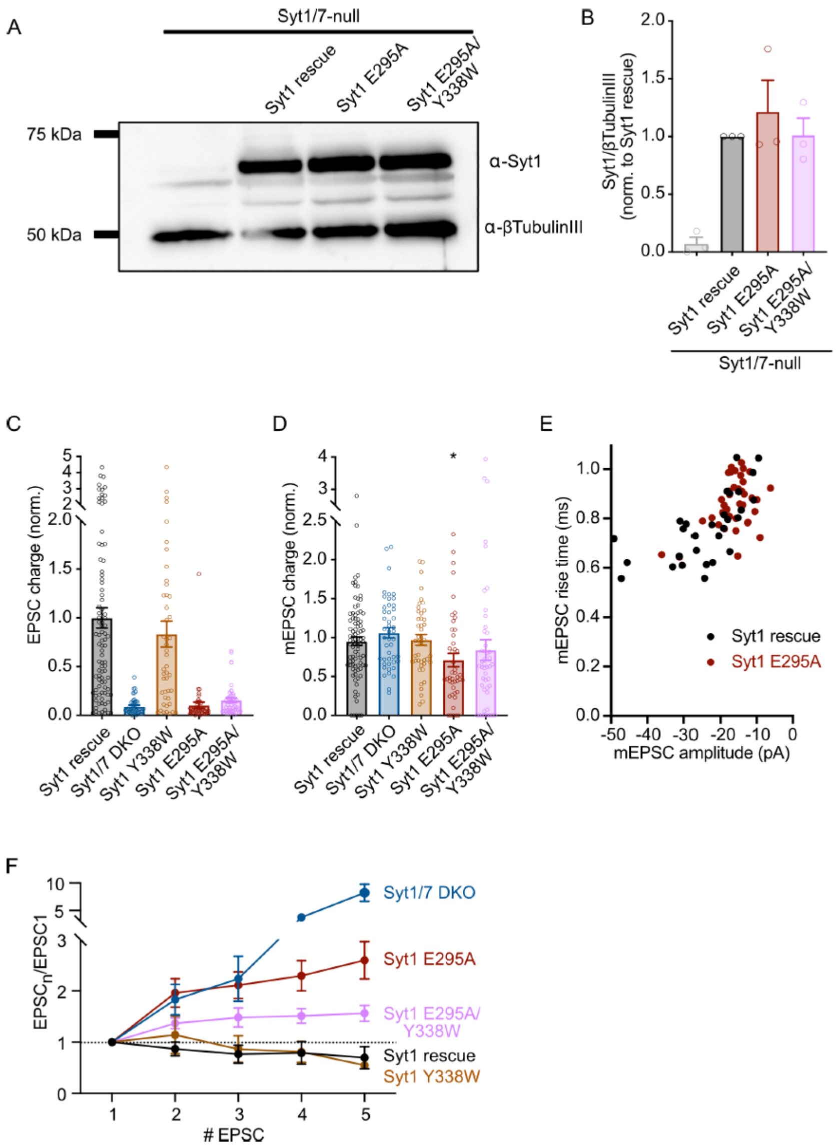


**Figure S5: Quantification of Syt1 mutant levels in hippocampal neurons and electrophysiological parameters of autaptic hippocampal mouse neurons rescued with Syt1 E295A and E295A/Y338W mutants from Region I.** (**A**) Example immunoblot and (**B**) quantification of protein levels for each experimental group. Proteins were detected using antibodies against beta-TubulinIII as loading control and Syt1 as the protein of interest. Protein expression for each mutant is normalized to Syt1 WT-rescue protein level. n = 3 replicates. (**C**) Quantification of the EPSC charged recorded for Syt1/7 DKO autaptic hippocampal neurons rescued with Syt1 WT, Syt1 E295A or Syt1 E295A/Y338W obtained from the same neurons as in (Fig. 5C). (**D**) Quantification of the miniature EPSC amplitude obtained from the same neurons as in (Fig. 5C). **(E)** Correlation plot between the mEPSC rise time and the mEPSC amplitude for a control group (Syt1 rescue) and Syt1 E95A mutant. The elevated rise time in the E295A mutant group is likely due to the detection algorithm, as the WT control shows a similar rise time to amplitude behavior. **(F)** Quantification of short-term plasticity measured by 5 stimulations at 50 Hz obtained from the same neurons as in (Fig. 5C). Between 44 and 45 neurons per group from 3 independent cultures were recorded and are shown as mean +/- SEM. ns: not significant, *p<0.05, **p<0.01, ***p<0.001 and ****p<0.0001.
